# *De novo* progesterone synthesis in plants

**DOI:** 10.1101/2023.07.19.549634

**Authors:** Rongsheng Li, Shuang Guo, Dong Wang, Tingting Yang, Xueli Zhang, Zhubo Dai

**Affiliations:** Tianjin Institute of Industrial Biotechnology, Chinese Academy of Sciences, Tianjin, 300308, China; University of Chinese Academy of Sciences, Beijing 100049, China

**Keywords:** Progesterone, C21 steroids, Biosynthetic pathway, Plants

## Abstract

The essential roles of progesterone and other C21 steroids in animals are well-documented. Progesterone is an essential hormone for females to maintain a regular menstrual cycle and pregnancy, while also exhibiting anti-inflammatory and antitumor effects. While the biosynthesis pathway of C21 steroids is comprehensively understood in animals, the synthesis mechanisms of progesterone in plants remain unclear. To our best knowledge, this is the first study to elucidate the complete pathway for progesterone biosynthesis in the plant *Marsdenia tenacissima*, involving the two sterol side chain cleaving cytochrome P450 enzymes (P450scc) Mt108 or Mt150, as well as the Δ^5^-3β-hydroxysteroid dehydrogenase/Δ^5^-Δ^4^ ketosteroid isomerase MtHSD5. This critical discovery paves the way for the sustainable synthesis of steroid hormone drugs using either plants or microbial host cells.

---

Progesterone is an essential hormone for females to maintain a regular menstrual cycle and pregnancy(Kolatorova et al., 2022). In addition, progesterone and other C21 steroids are widely used as drugs, constituting major anti-inflammatory, contraceptive, and anticancer agents(McMaster and Rays, 2008; Schleimer et al., 1981). Currently, the two principal industrial methods for synthesizing C21 steroids include chemical synthesis and microbiological bioconversion, which primarily use cholesterol or phytosterols as the starting material(Garcia et al., 2012; Velluz et al., 1960; Woodward et al., 1952). The third potential route, which is based on the development of synthetic biology and involves *de novo* biosynthesis, has great untapped potential(Ajikumar et al., 2010; Bai et al., 2016; Dusseaux et al., 2020; Gao et al., 2023; Liu et al., 2018; Ma et al., 2021; Peng et al., 2018; Srinivasan and Smolke, 2020; Zha et al., 2022; Zhang et al., 2022). Early studies attempted to biosynthesize C21 steroids such as progesterone and hydrocortisone from glucose, using pathways derived from animal sources(Szczebara et al., 2003). However, the low activity of relevant enzymes has limited the development of this route. Therefore, identifying efficient key enzymes such as the P450 sterol side chain cleaving enzyme (P450scc) is crucial to overcome this bottleneck.

In earlier studies, scientists have detected C21 steroids in plants at levels of up to 12% dry cell weight(Zheng et al., 2010). However, the biosynthetic pathway of these compounds and their catalytic activity is unknown(Meitinger et al., 2015; Zheng et al., 2014). In our work, a synthetic biology platform for elucidating the biosynthetic process of C21 steroids was established. This study identified, for the first time, the complete biosynthetic pathway of progesterone in *Marsdenia tenacissima*, a medicinal plant rich in C21 steroids (Figs. 1A and B). The pathway comprises three enzymes, the two P450scc Mt108 and Mt150, as well as the Δ^5^-3β hydroxysteroid dehydrogenase/Δ^5^-Δ^4^ ketosteroid isomerase MtHSD5 (Fig.1C). The elucidation of this pathway provides a crucial foundation for understanding *de novo* progesterone biosynthesis in plants, paving the way for the green synthesis of C21 steroid drugs using advanced cell factories in the future.

**Figure 1:**
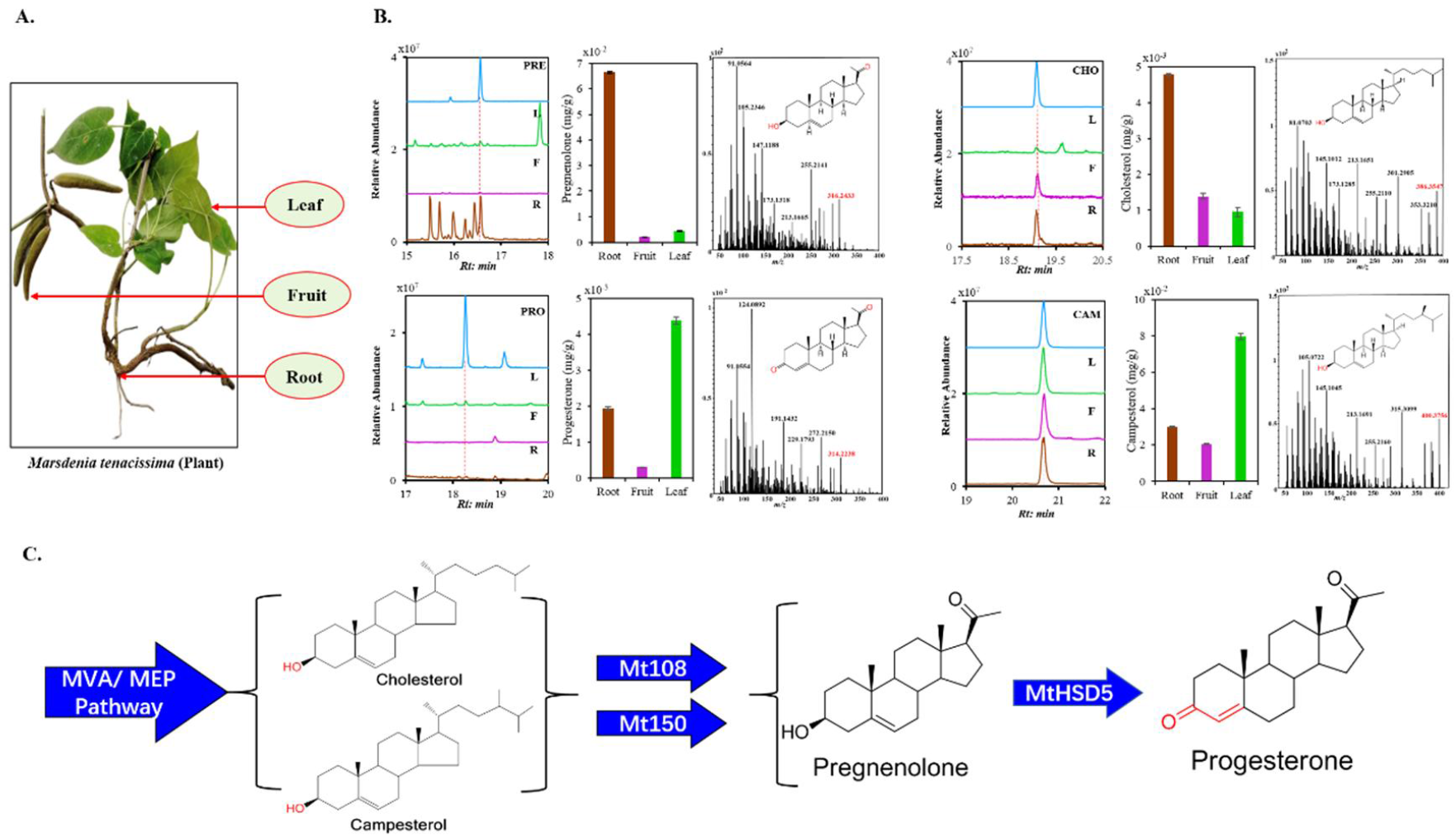
The distribution of progesterone and its possible precursors in *Marsdenia tenacissima*. A. Plant material of *Marsdenia tenacissima*; B. The relative concentrations of progesterone and its precursors in various tissues; C. The proposed biosynthesis of progesterone in *M. tenacissima*. Single arrows represent one-step conversions. The Mt108 and Mt150 genes of *M. tenacissima* encode a P450 sterol side chain cleaving enzyme (P450scc). The MtHSD5 gene of *M. tenacissima* encodes Δ^5^-3β hydroxysteroid dehydrogenase /Δ^5^-Δ^4^ ketosteroid isomerase.

## Results and Discussion

### 1. The distribution of progesterone and its possible precursors in plants

Although progesterone is mainly a hormone produced by animals, it is found alongside other C21 steroids in a number of plant species from the families Apocynaceae and Asclepiadaceae, including *M. tenacissima, Periploca sepium, Cynanchum caudatum*, and *Digitalis purpurea*(Gasic et al., 2023; Zheng et al., 2014). In particular, *M. tenacissima* can contain up to 12% cell dry weight of these compounds(Zheng et al., 2010). However, little was known about the distribution of progesterone, as well as its possible precursors such as cholesterol, campesterol and pregnenolone, in these species.

In order to investigate the distribution of progesterone, along with its potential biosynthetic precursors including cholesterol, campesterol and pregnenolone, various plant tissues were collected from *M. tenacissima*, including roots, leaves, and fruits (Fig. 1A). Quantitative GC-MS analysis revealed the presence of progesterone in all the examined tissues, with the leaves showing the highest concentrations, reaching up to 4.3 μg/g (Fig. 1B). Pregnenolone, the corresponding precursor, primarily accumulated in the roots, with a concentration of approximately 66.5 μg/g (Fig. 1B). Additionally, common phytosterols such as cholesterol and campesterol accumulated in the roots, leaves, and fruits. Cholesterol primarily accumulated in the roots at approximately 4.8 μg/g, whereas campesterol accumulated in the leaves at around 79.6 μg/g (Fig. 1B).

### 2. Construction of chassis strains for screening cytochrome P450 enzymes

Recently, the use of synthetic biology platforms for functional gene identification has been established as a workable strategy, and has been successfully applied for investigating the biosynthesis of saponins and triterpenoids(Dai et al., 2019a; Li et al., 2021b; Wang et al., 2020).

Progesterone is derived from cholesterol through the action of P_SCC_ enzyme in animals(Szczebara et al., 2003). To determine whether a similar pathway exists in plants, we first engineered a chassis strain capable of producing both cholesterol and phytosterols. In the chassis strain *S. cerevisiae* YSBYT5(Shi et al., 2021), which is an engineered strain with an optimized mevalonate (MVA) pathway, we deleted the gene cluster of gal1, gal7, and gal10 to obtain strain BY-RS01, which does not metabolize galactose but can still use it as an inducer(Paddon et al., 2013). We respectively placed the HMGR enzyme genes from three different sources under the control of the GAL1, GAL7, and GAL10 promoters, and integrated the resulting cassettes into the chromosomal ypl062w locus of *S. cerevisiae* strain BY-RS01, resulting in strain BY-RS02. Subsequently, the cytochrome P450 reductase gene *VvCPR* from grapes (*Vitis vinifera*)(Dai et al., 2019a), sterol Δ^7^-reductase gene *StDWF5* from potatoes (*Solanum tuberosum*)(Xu et al., 2022), and Δ^24^-dehydrocholesterol reductase gene *GgDHCR24* from domestic chicken (*Gallus gallus*) (Xu et al., 2022)were inserted into the *atf2* site of BY-RS02 to obtain the engineered cholesterol-producing strain CHOL01. The *S. cerevisiae* strain CHOL01 was obtained by integrating the yeast protein disulfide isomerase (PDI1)(Noiva and Lennarz, 1992) chaperone gene and the endoplasmic reticulum membrane formation gene *INO2* (Kim et al., 2019) into the *HO* site of the cholesterol-producing strain CHOL01, resulting in strain CHOL02. This resulted in the creation of the CHOL02 chassis strain capable of synthesizing 39.4 mg/L cholesterol (Fig. 2A).

**Figure 2:**
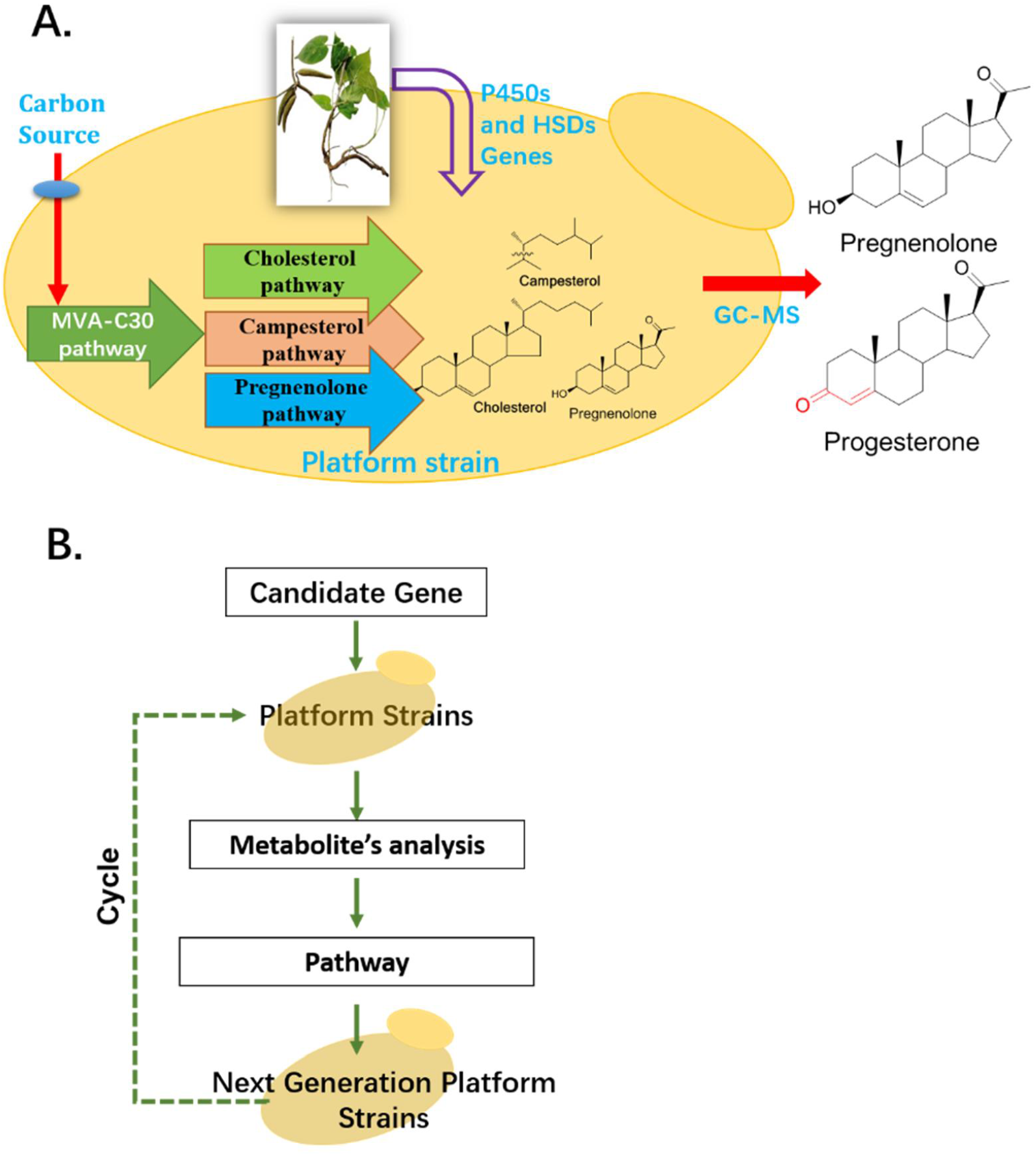
The synthetic biology platform used to elucidate the complete biosynthetic pathway of PNPs. A. Elucidation of the complete biosynthetic pathway of the progesterone in *M. tenacissima* using a synthetic biology platform; B. Technical flow chart of the platform.

Additionally, during the construction of the chassis strain CAOL01 for campesterol production, the integration of the *VvCPR* and *StDWF5* genes into the *atf2* locus of engineered strain BY-RS02 led to a campesterol yield of 49.3 mg/L (Fig.2A).

### 3. Functional analysis of the MT108 and Mt150 genes

Based on the distribution analysis of C21 steroid metabolites in plants (Fig.1B), transcriptome sequencing was performed on the roots, fruits, and leaves of *M. tenacissima*. Next, we used the full-length amino acid sequences of the 151 contigs, together with the amino acid sequences of representative CYPs involved in steroid biosynthesis(Christ et al., 2019; Malhotra and Franke, 2022), to conduct a phylogenetic analysis. A total of 7 CYP genes were selected based on the phylogeny of the corresponding CYP proteins for subsequent cloning (Fig.3, Marked with red squares). Finally, the candidate CYPs were successfully cloned and inserted into the high-copy plasmid pRS425 for functional gene characterization.

**Figure 3:**
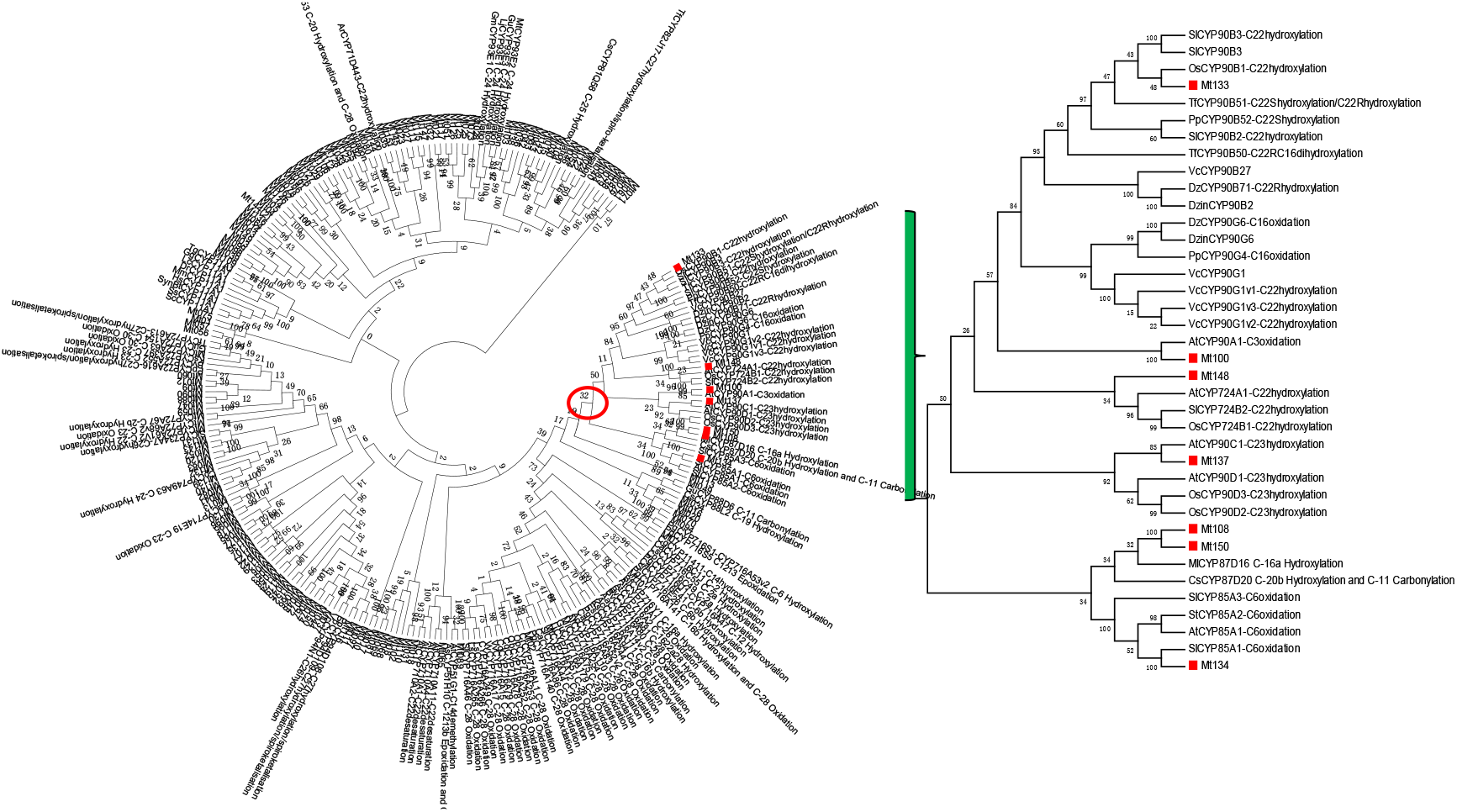
A phylogenetic tree of CYP enzymes. The red squares indicate the candidate CYPs from *M. tenacissima*. The evolutionary history was inferred using the Neighbor-Joining method. The amino acids were aligned using CLUSTALW. Bootstrap values from 1000 retrials are indicated at each branch.

To identify the enzymes responsible for the biosynthesis of progesterone, we utilized the CYP gene from *M. tenacissima* to screen the target genes using a “plug- and-play” workflow (Fig. 2B). We identified two genes, MT108 and Mt150, whose encoded proteins exhibited activity consistent with the sterol side chain cleaving enzyme (Fig. 4). To further evaluate the P450 activity, the Mt108 and Mt150 genes were integrated into the *yjl064w* site of the campesterol-producing strain CAOL01 under the control of the GAL1 promoter. GC-MS analysis of strains CAOL01-Mt150 and CAOL01-Mt108 expressing the cytochrome P450 enzyme genes confirmed the production of pregnenolone by Mt108 or Mt150, as validated by a comparison with an authentic reference standard (Fig. 5).

**Figure 4:**
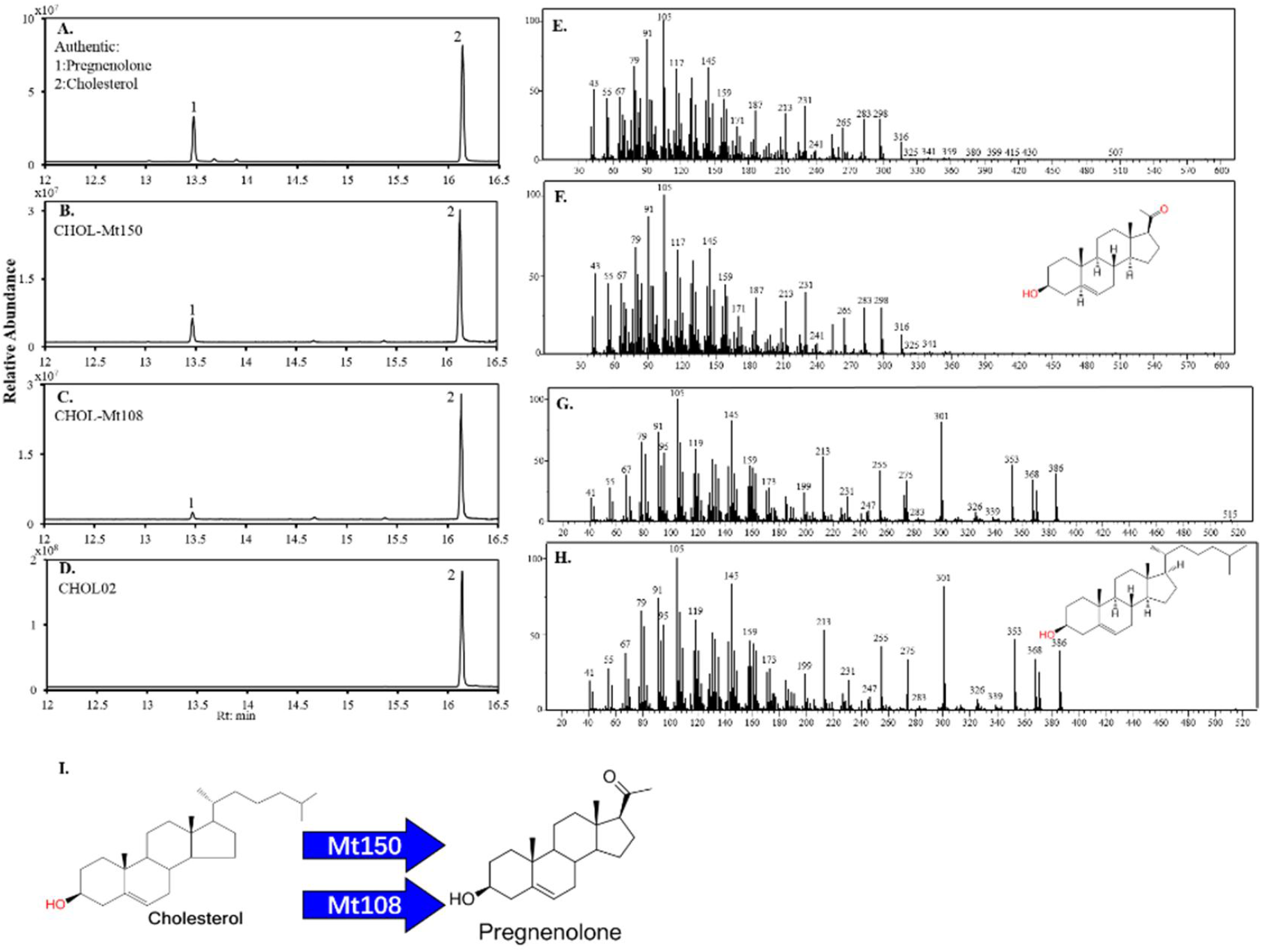
Identification of the key enzyme involved in the biosynthesis of pregnenolone from cholesterol in plants. A shows the GC-MS chromatogram of the pregnenolone and cholesterol standards, with peaks 1 and 2 corresponding to pregnenolone and cholesterol, respectively; B, C, and D are the GC-MS chromatograms of fermentation products from strains CHOL-Mt150, CHOL-Mt108, and CHOL02, respectively; E and G are the MS spectra of peaks 1 and 2 among the fermentation products of strains CHOL02, CHOL-Mt150, and CHOL-Mt108, respectively; F and H are the MS spectra of pregnenolone and cholesterol, respectively; I shows the proposed biosynthetic pathway of pregnenolone synthesis from cholesterol in plants.

**Figure 5:**
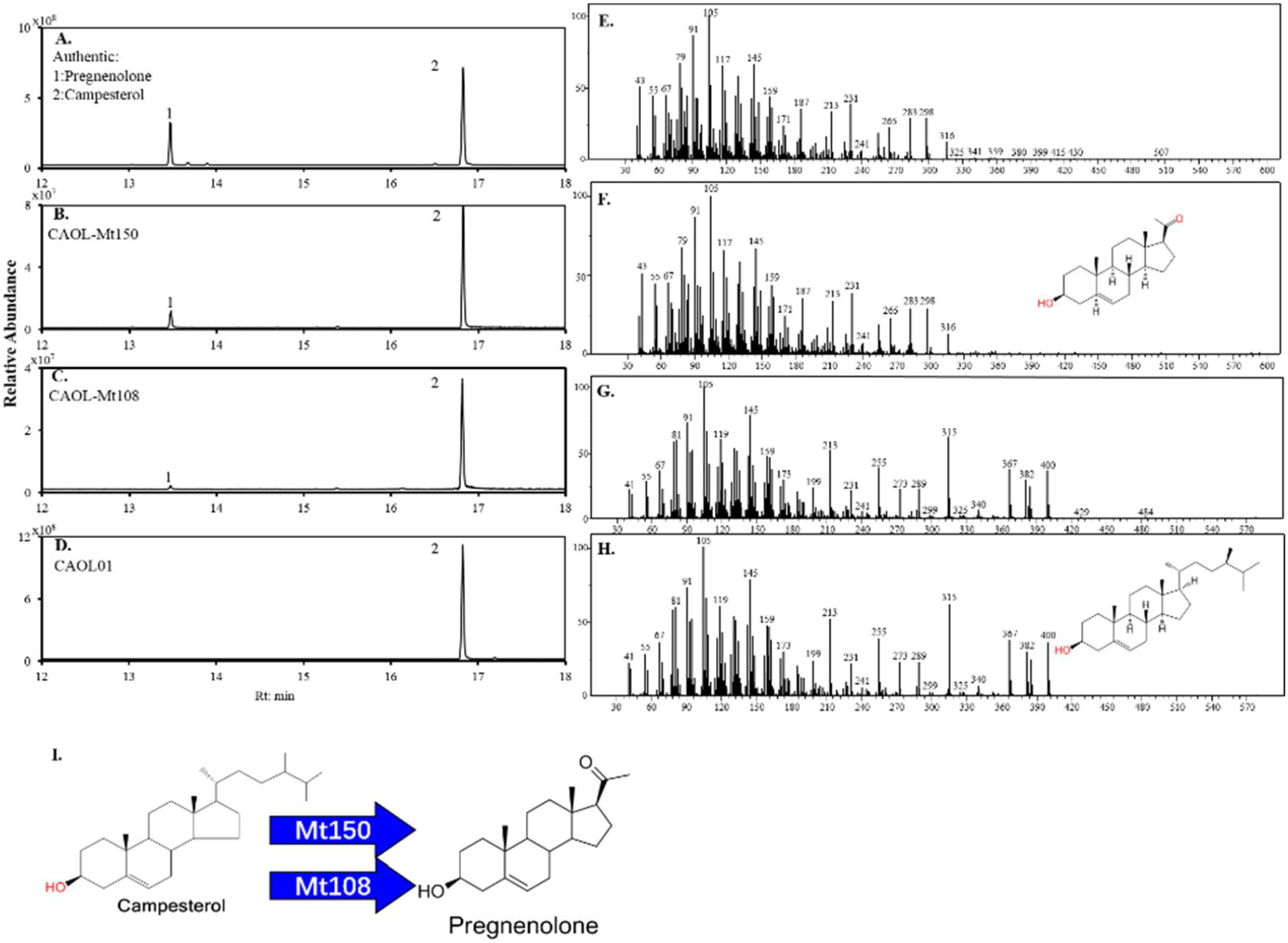
Identification of the key enzyme involved in the biosynthesis of pregnenolone from campesterol in plants. A shows the GC-MS peaks for the pregnenolone and campesterol reference standards, with peaks 1 and 2 corresponding to pregnenolone and campesterol, respectively; B, C, and D are the GC-MS chromatograms of the fermentation products of strains CAOL-Mt150, CAOL-Mt108, and CAOL01, respectively; E and G are the mass spectra of peaks 1 and 2 among the fermentation products of strains CAOL-Mt150, CAOL-Mt108, and CAOL01, respectively; F and H are the mass spectra of pregnenolone and campesterol, respectively; I shows the proposed biosynthetic pathway of pregnenolone synthesis from campesterol in plants.

### 4. Discovery of the enzyme catalyzing the conversion of pregnenolone into progesterone

Previous studies indicated the possibility that both Δ5-3β-hydroxysteroid dehydrogenase (3β-HSD) and Δ5-Δ4 ketosteroid isomerase (KSI) participate in the conversion of pregnenolone into progesterone in plants(Herl et al., 2006; Meitinger et al., 2015). To further identify the functional genes encoding enzymes responsible for catalyzing the conversion of pregnenolone into progesterone in plants, the engineered strain PG-Mt108 was constructed to produce progesterone by overexpressing MtHSD5 in the pregnenolone-producing strain. The engineered strain successfully produced detectable amounts of progesterone as confirmed by GC-MS analysis, successfully completing the pathway (Fig. 6). These results indicated that MtHSD5 is a multifunctional enzyme that possesses both Δ^5^-3β-hydroxysteroid dehydrogenase and Δ5-Δ4 ketosteroid isomerase activities.

**Figure 6:**
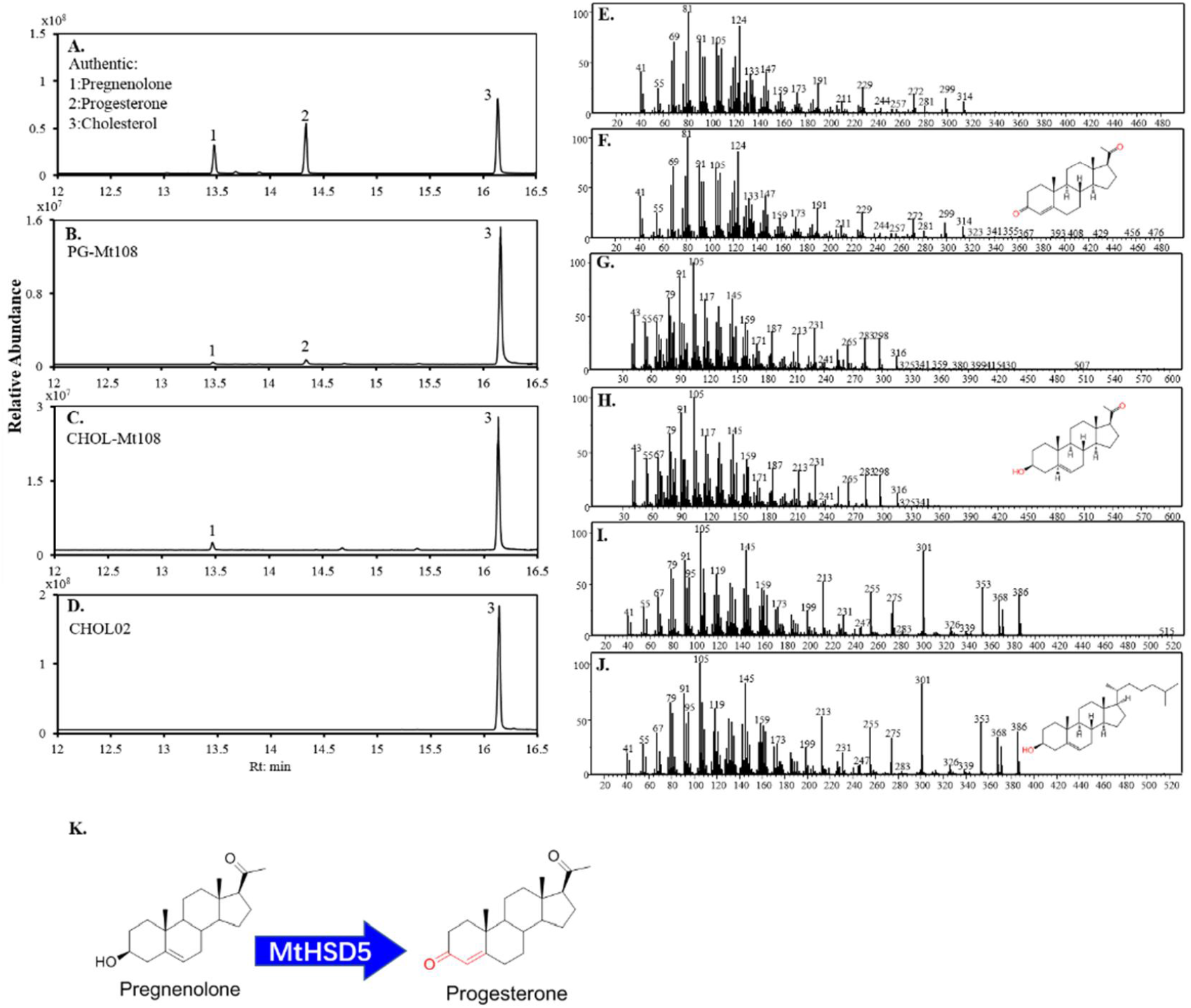
Identification of the key enzyme involved in the biosynthesis of progesterone from pregnenolone in plants. A shows the GC-MS peaks of the authentic reference standards, with peaks 1, 2, and 3 corresponding to pregnenolone, progesterone, and cholesterol, respectively; B, C, and D are the GC-MS chromatograms of the fermentation products of strains PG-Mt108, CHOL-Mt108, and CHOL02, respectively; E, G, and I are the mass spectra corresponding to peaks 1, 2, and 3 among the fermentation products of strains CHOL02, CHOL-Mt108, and PG-Mt108, respectively; F, H, and J are the mass spectra of progesterone, pregnenolone, and cholesterol, respectively; K represents the proposed pathway of progesterone synthesis from pregnenolone in plants.

In conclusion, we have successfully identified the complete biosynthetic pathway of progesterone in plants, providing a solid foundation for understanding the *de novo* synthesis of steroid hormones in plants, with great potential for eco-friendly drug synthesis in the future.

## Materials and methods

### Strains and culture conditions

The *S. cerevisiae* platform strains CHOL01, CHOL02, CAOL01, BY-RS01 and BY-RS02 were cultured at 30 °C in SD medium with 20 g/L glucose, lacking tryptophan, leucine, histidine and uracil, when appropriate. SD-Trp-Ura (synthetic complete drop-out medium with 2% glucose, without tryptophan and uracil) was used for the selection of yeast strains carrying both the Cas9 plasmid 43802 and guide RNA (gRNA) plasmid 43803. The 5-FOA-SD-Trp medium (SD-Trp medium with 1 mg/mL 5-fluoroorotic acid) was used to remove the gRNA expression plasmid. All cloning procedures were carried out in *Escherichia coli* T1 (TransGen Biotech, Beijing, China), which was grown at 37 °C in LB medium supplemented with 100 mg/mL ampicillin when appropriate.

### RNA-Seq and gene cloning

*M. tenacissima* plants were collected near the city of Honghe in Yunnan Province, China. The RNA from the roots, fruits and leaves of *M. tenacissima* was isolated using TRIzol reagent. A total of 259.6 million paired-end raw reads were assembled *de novo* using Trinity software (v2.4.0). The candidate full-length CYP sequence was confirmed by pfam (PF00067), and the candidate full-length HSD sequence was confirmed by pfam (PF01073) prior to cloning.

### Plasmid construction

*M. tenacissima* plantlets were induced with 1 mM MeJA for 24 h, after which the RNA was isolated using the TRIzol method, and the cDNA was obtained using the PrimeScript first Strand cDNA Synthesis Kit (Takara, Dalian, China). The open reading frames of genes were amplified by PCR from the cDNA of *M. tenacissima* and ligated into the vector using the seamless cloning kit (Beyotime Biotech, Shanghai, China). The primer sequences are summarized in Table 3. All plasmids used in this study are listed in Table 1.

**Table 1.**
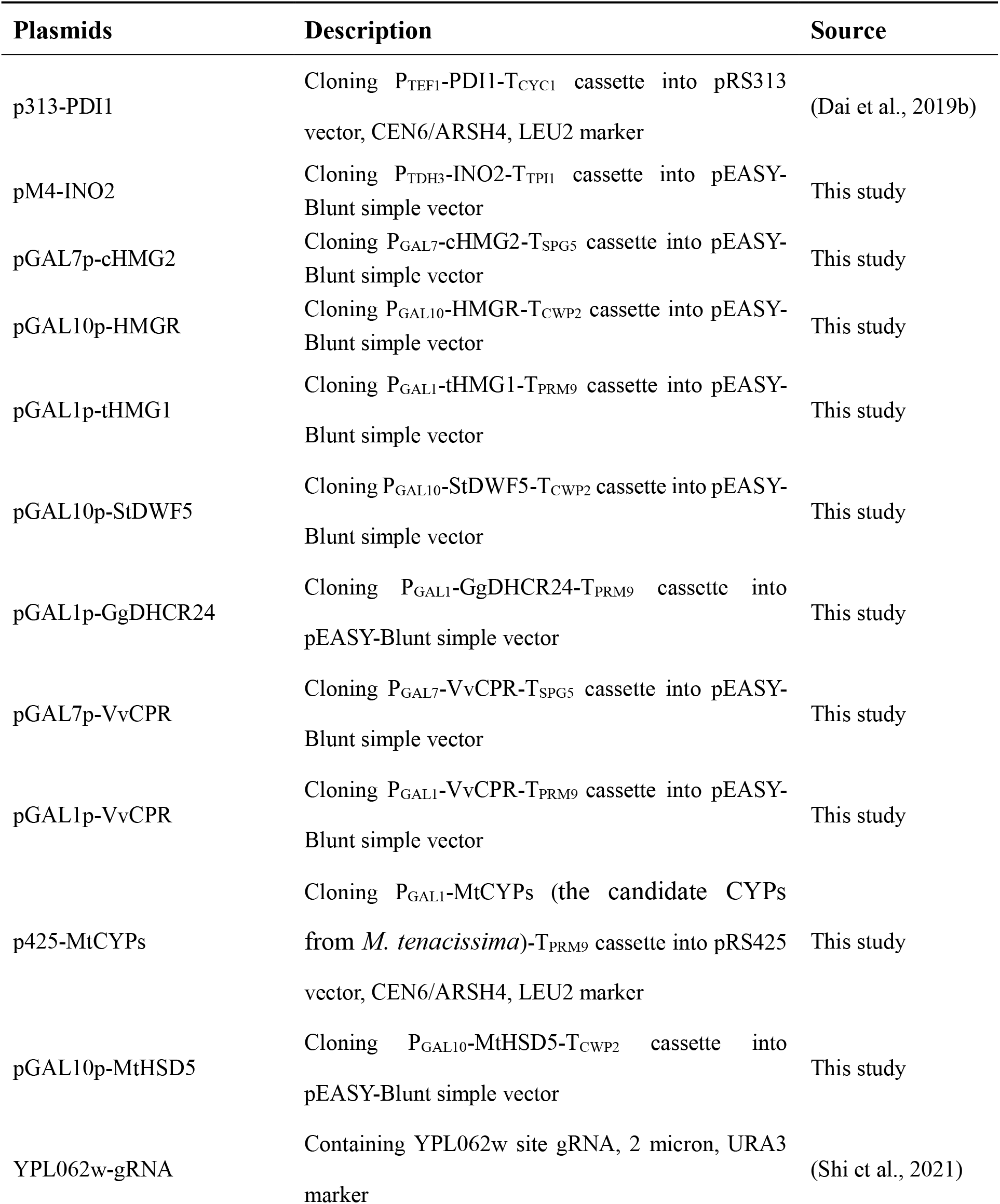
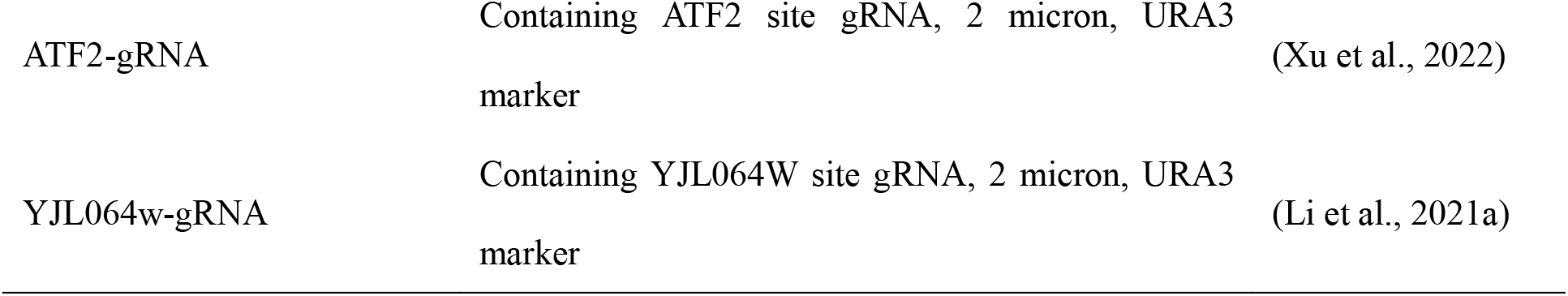
Plasmids used in this study.

**Table 2.** *S. cerevisiae* strains used in this study.

| Strains | Description | Source |
| --- | --- | --- |
| YSBYT5 | BY4742, *trp1Δ*, δDNA:: P<sub>PGK1</sub>-*tHMG1*-T<sub>ADH1</sub>-P<sub>TEF1</sub>-*LYS2*-T<sub>CYC1</sub>, *leu2*:: P<sub>PGK1</sub>-*tHMG1*-T<sub>ADH1</sub>-P<sub>PDC1</sub>-*ERG12*-T<sub>ADH2</sub>-P<sub>ENO2</sub>-*IDII*-T<sub>PDC1</sub>-P<sub>PYK1</sub>-*ERG19*-T<sub>PGI1</sub>-P<sub>TEF2</sub>-*HMGR*-T<sub>ENO2</sub>-P<sub>FBA1</sub>-*ERG13*-T<sub>TDH2</sub>-P<sub>TDH3</sub>-*ERG8*-T<sub>TPI1</sub>-P<sub>TEF1</sub>-*ERG10*-T<sub>CYC1</sub> | (Shi et al., 2021) |
| BY-RS01 | YSBYT5, *gal1Δ*, *gal7Δ*, *gal10Δ* | This study |
| BY-RS02 | BYRS01, *yp1062w*:: T<sub>SPG5</sub>-*cHMG2*-P<sub>GAL7</sub>-T<sub>CWP2</sub>-*HMGR*-P<sub>GAL10-GAL1</sub>-*tHMG1*-T<sub>PRM9</sub> | This study |
| CHOL01 | BYRS02, *atf2*:: T<sub>SPG5</sub>-*VvCPR*-P<sub>GAL7</sub>-T<sub>CWP2</sub>-*StDWF5*-P<sub>GAL10-GAL1</sub>-*GgDHCR24*-T<sub>PRM9</sub> | This study |
| CHOL02 | CHOL01, *ho*:: P<sub>PGK1</sub>-*tHMG1*-T<sub>ADH1</sub>-P<sub>TDH3</sub>-*INO2*-T<sub>TPI1</sub>-P<sub>TEF1</sub>-*PDII*-T<sub>CYC1</sub> | This study |
| CAOL01 | BYRS02, *atf2*:: T<sub>CWP2</sub>-*StDWF5*-P<sub>GAL10-GAL1</sub>-*VvCPR*-T<sub>PRM9</sub> | This study |
| CHOL-MtCYPs | CHOL02, *yjl064w*:: P<sub>GAL1</sub>-*MtCYPs*-T<sub>PRM9</sub> | This study |
| CHOL-Mt108 | CHOL02, *yjl064w*:: P<sub>GAL1</sub>-*MtCYP108*-T<sub>PRM9</sub> | This study |
| CHOL-Mt150 | CHOL02, *yjl064w*:: P<sub>GAL1</sub>-*MtCYP150*-T<sub>PRM9</sub> | This study |
| CAOL-Mt108 | CAOL01, *yjl064w*:: P<sub>GAL1</sub>-*MtCYP108*-T<sub>PRM9</sub> | This study |
| CAOL-Mt150 | CAOL01, *yjl064w*:: P<sub>GAL1</sub>-*MtCYP150*-T<sub>PRM9</sub> | This study |
| PG-Mt108 | CHOL02, *yjl064w*:: T<sub>CWP2</sub>-*MtHSD5*-P<sub>GAL10-GAL1</sub>-*MtCYP108*-T<sub>PRM9</sub> | This study |

**Table 3.**
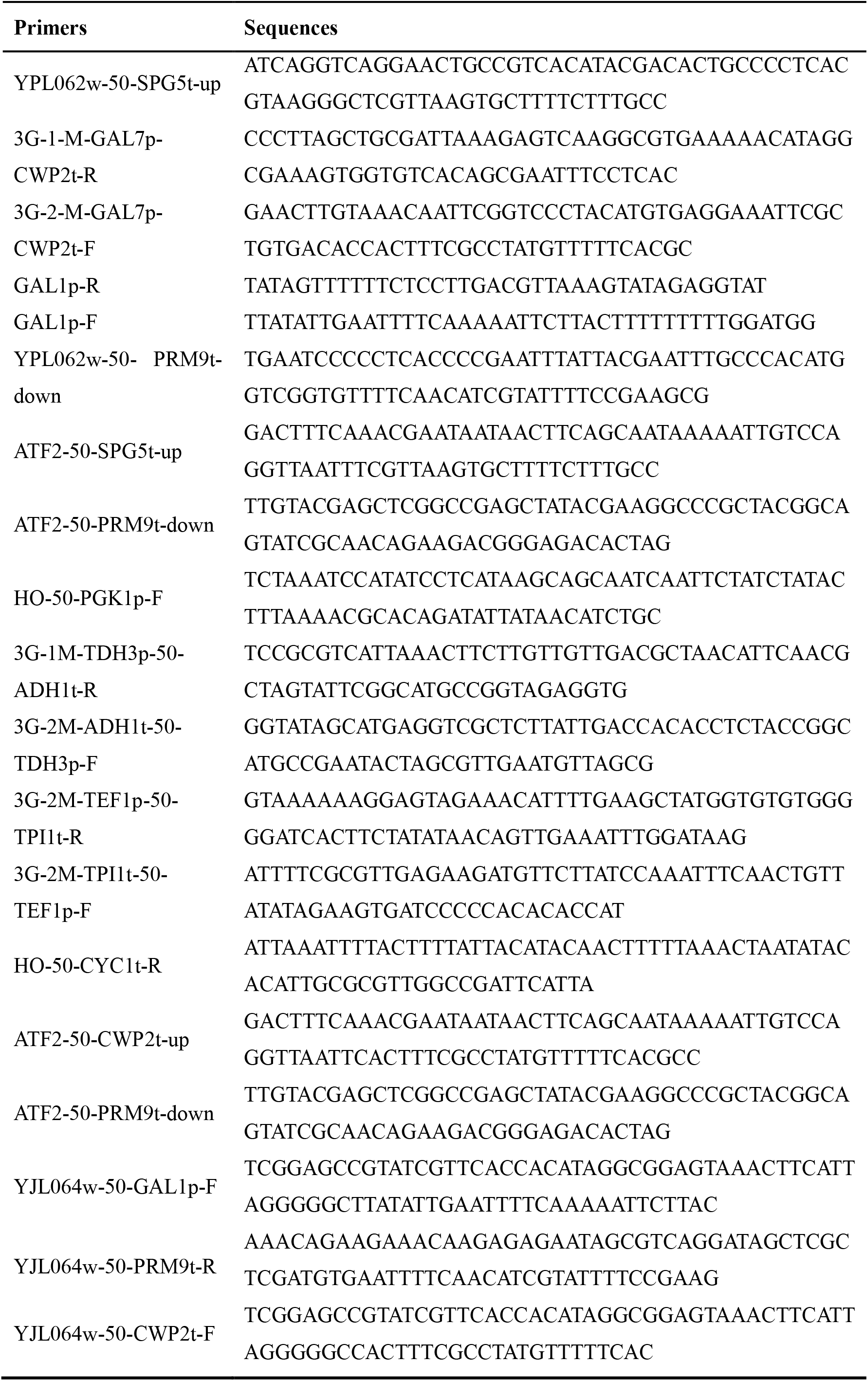
Primers used in this study.

### Strain construction

Transformation of yeast strains was performed using a conventional electroporation method as described previously (Dai et al., 2013). All the strains used in this study are listed in Table 2.

### Shake-Flask Cultivation

Yeast strains were cultured overnight in uracil and tryptophan dropout medium with 2% glucose. Then, the seed culture was used to inoculate 15 ml of the same medium (5% inoculum), followed by culture at 30 ºC and 250 rpm for 48 h. The culture was placed in a sterile 50ml centrifuge tube, centrifuged at 6000 rpm for 5 min and the supernatant discarded. Then, the cells were resuspended in 15ml of uracil and tryptophan dropout induction medium (1% galactose, 2% ethanol), followed by further fermentation at 30 ºC and 250 rpm for 72 h.

## Chemical analysis

### Analysis of steroid compounds in plant tissues of *M. tenacissima*

Samples comprising 10 g of the dried roots, leaves, and fruits of *M. tenacissima* were weighed and soaked in 50 ml of 70% ethanol overnight, followed by ultrasonic extraction for 2h two times. The extracts were collected and evaporated at reduced pressure, after which 10 ml of acetone/methanol (1:1, v/v) was added to fully dissolve the extract. Subsequently, the organic phase was collected, and 1 μL was subjected to combined gas chromatography-mass spectrometry (GC-MS) analysis on a 7890A GC system, equipped with a 200 Accurate-Mass selective detector and an DB-35ms GC column (30 m × 0.25 mm, 0.5-μm film thickness). The temperature program encompassed 50 ^°^C for 1 min, ramp to 300 ^°^C (20 ^°^C/min) for 12.5 min and a final constant hold at 300 ^°^C for 30 min. Authentic reference standards of pregnenolone (≥98%), progesterone (≥98%), cholesterol (≥99%), and campesterol (≥98%), were purchased from Yuanye Bio-Technology Co., Ltd (Shanghai, China).

### Analysis of steroid compounds in recombinant yeast strains

Sample preparation was done as described previously(Xu et al., 2022). The cell pellets were harvested, disrupted, and extracted with 1mL of acetone/methanol (1:1, v/v), after which 1 μL was subjected to combined gas chromatography-mass spectrometry (GC-MS) analysis on a Thermo Scientific Trace1300 GC system, equipped with a 200 Accurate-Mass selective detector and an DB-35ms GC column (30 m × 0.25 mm, 0.5-μm film thickness). The temperature program encompassed 100 °C for 1 min, ramp to 300 °C (20 °C/min) for 10 min and a final constant hold at 300 °C for 15 min.

## Acknowledgments

This research was supported by National Key Research and Development Project (2020YFA0908000, 2019YFA0905300). Dr. Zhubo Dai was supported by Youth Innovation Promotion Association of CAS (2015138). We thank Dr. Guodong Li (Yunnan University of Chinese Medicine) for providing the plant material of *Marsdenia tenacissima*.

## Additional information

This work has been included in patent applications by the Tianjin Institute of Industrial Biotechnology

## Notes

### Competing Interest Statement

The authors have declared no competing interest.

